# Transfer of mitochondrial DNA into the nuclear genome during gene editing

**DOI:** 10.1101/2023.07.19.549443

**Authors:** Jinchun Wu, Yang Liu, Liqiong Ou, Tingting Gan, Zhengrong Zhangding, Shaopeng Yuan, Mengzhu Liu, Xinyi Liu, Jiasheng Li, Jianhang Yin, Changchang Xin, Ye Tian, Jiazhi Hu

**Affiliations:** The MOE Key Laboratory of Cell Proliferation and Differentiation, School of Life Sciences, Center for Life Sciences, Genome Editing Research Center, Peking University, Beijing 100871, China; State Key Laboratory of Molecular Developmental Biology, Institute of Genetics and Developmental Biology, Chinese Academy of Sciences

**Keywords:** Genome editing, DdCBE, mitochondrial base editor, CRISPR-Cas9, mitochondrial integration into nuclear DNA, TREX1/2

## Abstract

Mitochondria serve as the cellular powerhouse, and their distinct DNA makes them a prospective target for gene editing to treat genetic disorders. However, the impact of genome editing on mitochondrial DNA (mtDNA) stability remains a mystery. Our study reveals previously unknown risks of genome editing that both nuclear and mitochondrial editing cause broad transfer of mitochondrial DNA segments into the nuclear genome in various cell types including human cell lines, primary T cells, retinal cells, and mouse embryos. Furthermore, drug-induced mitochondrial stresses and mtDNA breaks exacerbate this transfer of mtDNA into the nuclear genome. Notably, we observe that the newly developed mitochondrial base editor DdCBE can also cause widespread mtDNA integrations. However, we provide a practical solution to suppress the transfer of mtDNA by co-expressing TREX1 or TREX2 exonucleases during DdCBE editing. These findings also shed light on the origins of mitochondrial-nuclear DNA segments.

## Introduction

The human genome consists of two copies of nuclear DNA and a few hundred to a few thousand copies of mitochondrial DNA (mtDNA), which are separated by the nuclear and transferred to the nuclear genome during the endosymbiotic evolution (Gray et al., 1999). Consequently, the human genome contains hundreds of nuclear-mitochondrial DNA segments (NUMTs) and still undergoes the natural transfer of mtDNA to the nuclear DNA (Wei et al., 2020; Wei et al., 2022a; Ju et al., 2015). *De novo* NUMTs were detected in approximately one per 10^4^ birth and 10^3^ cancers (Ju et al., 2015; Wei et al., 2022a). Meanwhile, NUMTs in tumor cells tend to be embedded in gene-rich regions of nuclear DNA, suggesting that they may be the causal integration of certain tumors (Wei et al., 2022a; Kopinski et al., 2021).

Recently, the focus of gene editing has extended from the nuclear DNA (Jiang and Doudna, 2017; Hsu et al., 2014; Anzalone et al., 2020; Liu et al., 2021) to mtDNA (Silva-Pinheiro and Minczuk, 2022; Barrera-Paez and Moraes, 2022), leading to the development of a series of highly effective mitochondrial editors. These tools aim to eliminate mutated mtDNA by inducing DNA double-stranded breaks (DSBs) with mitochondrial-targeted transcription activator-like effector nucleases (mitoTALEN) (Bacman et al., 2013; Bacman et al., 2018) or correcting mutations with mitochondrial base editors (Mok et al., 2020; Cho et al., 2022). DNA insertions of genomic or viral origins have been widely observed during gene editing exerted by TALEN or clustered regularly interspaced short palindromic repeats associated system (CRISPR-Cas) (Wu et al., 2022; Yin and Hu, 2022; Kosicki et al., 2018). However, it remains to be investigated whether mitochondrial or nuclear editing can result in the integration of mtDNA segments into the nuclear genome.

Here we found that genome editing on the nuclear DNA caused mtDNA-nuclear DNA fusion at the CRISPR-Cas9 target sites *in vitro* and *in vivo*, validated by both primer-extension-mediated sequencing (PEM-seq) and target sequencing. Moreover, high-fidelity Cas9 variants and base editors were unable to eliminate the transfer of mtDNA to nuclear DNA. Mitochondrial stresses increased the level of mtDNA-nuclear DNA fusion, and mitochondrial editing tools, such as mitoTALEN and DddA-derived cytosine base editor (DdCBE), also caused mtDNA fragility and resulted in the transfer of mtDNA to nuclear DNA. Finally, we found that co-expression of either TREX1 or TREX2 exonuclease in the mitochondria could be a plausible solution to suppress mtDNA fusion to the nuclear DNA during DdCBE treatment.

## Results

### Mitochondrial DNA fuses to CRISPR-Cas targeting sites in the nuclear genome

To investigate the integration of mtDNA segments into nuclear CRISPR-Cas target sites, we examined potential mtDNA sequences in the CRISPR-Cas9 editing products profiled by PEM-seq (Fig. 1A). PEM-seq utilizes a primer adjacent to the “bait” target site to capture “prey” sequence fused to the target DNA breaks through a one-round primer extension, rather than PCR. Subsequently, the single-stranded DNA products were isolated and subjected to library preparation (Fig. 1A)(Wu et al., 2022; Liu et al., 2022). In order to preclude potential errors in primer annealing, library preparation and sequencing, we conducted parallel PEM-seq analysis at each locus using genomic DNA from both edited and unedited cells (Fig. 1A). By employing PEM-seq, we successfully identified fusion junctions between mtDNA and the CRISPR target sites for all the tested editing loci, including *CLIC4*, *KLHL29*, *NLRC4*, *COL8A1*, *NUDT16*, *LNX1*, *FGF18*, *VEGFA_1/2*, *HBB*, *IFNγ*, and *P2RX5-TAX1BP3* in HEK293T cells, edited with CRISPR-Cas family editors including *Lb*Cas12a, *As*Cas12a, *Un1*Cas12f, CasMINI, and CasMINI_ge4.1 (Fig. 1B, Supplementary Table 1 and 2) (Xin et al., 2022). Since the human nuclear genome contains multiple embedded mtDNA segments, we only counted the chimeric reads where the prey sequences were better aligned to the mtDNA as mitochondria-nuclear fusions or translocations, exemplified by two fragments fused to *VEGFA_1* or *NLRC4* baits (Fig. S1A). The frequency of mtDNA integration at CRISPR-Cas target site varied from one per 10^3^ to 10^5^ edited events in HEK293T cells, with the Cas12 family exhibiting slightly lower levels of mtDNA integrations in comparison to *Sp*Cas9 (Fig. 1B-E and S1B-D). In contrast, unedited samples showed no mtDNA-nuclear DNA fusions for 10 of the 12 tested loci. Only two editing events out of 625,886 for *VEGFA_1* and one editing event out of 897,959 for *HBB* loci were detected in the unedited samples (Fig. 1B, Supplementary Table 1). However, these events contained the exact same sequences as those observed in the corresponding editing samples (Supplementary Table 1), indicating that the rare events in the control libraries may originate from clustering errors during Illumina sequencing.

**Figure 1.**
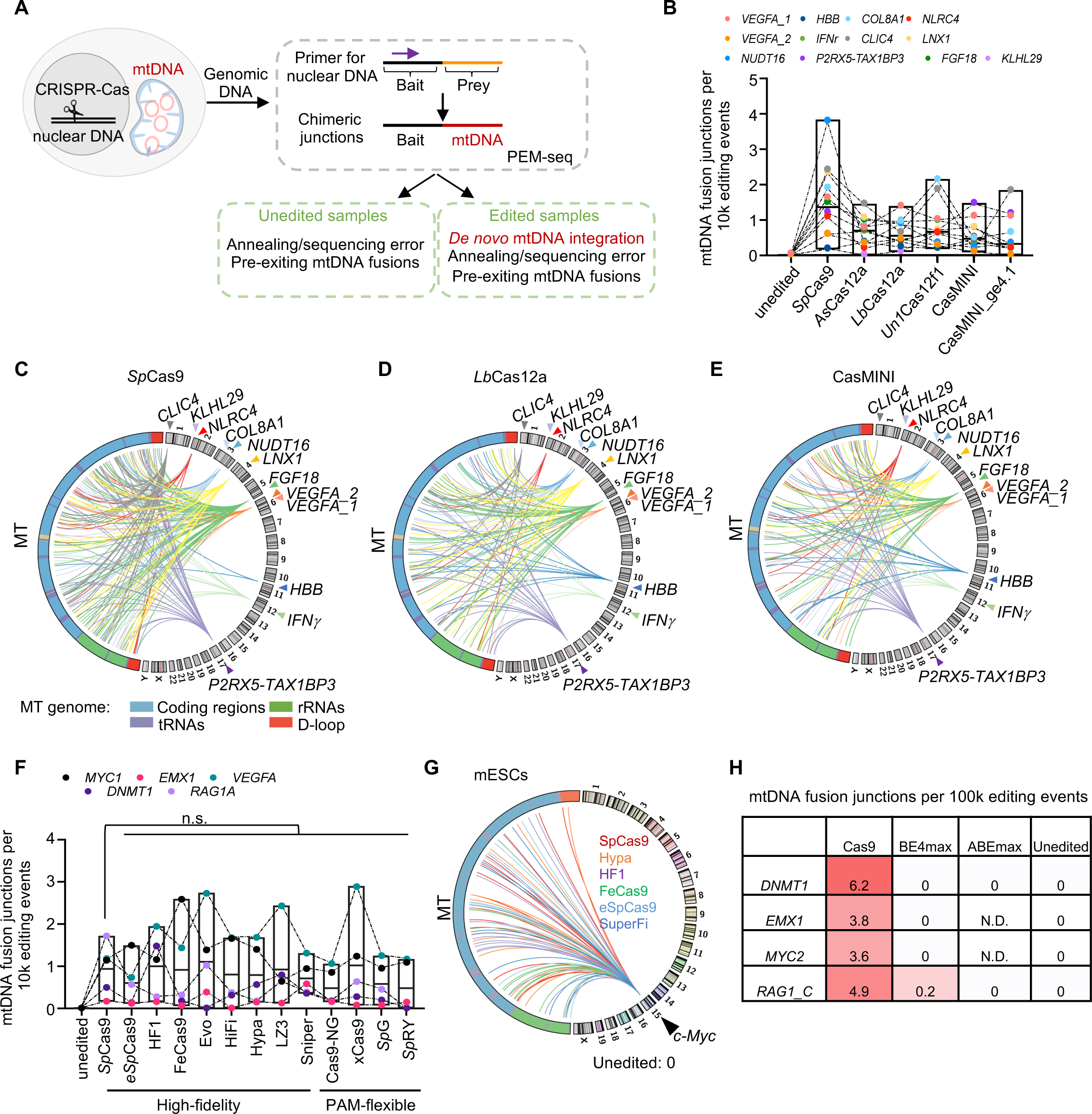
Identification of mtDNA fusion to nuclear target sites of CRISPR-Cas system. A. Schematic of mtDNA-nuclear DNA fusion captured by PEM-seq. The primer (purple arrow) located adjacent to the CRISPR-Cas9 target site (scissor) on the nuclear DNA is used to clone editing products (orange line), and the chimeric reads harboring nuclear DNA around the editing site (black line) and mtDNA (red line), with MAPQ >30, were identified as mtDNA-nuclear DNA fusion. B. Box plot showing the frequency of mtDNA-nuclear DNA fusion events out of editing events at CRISPR-Cas target sites (colorful dots) under editing of CRISPR-Cas enzymes. C. Circos plot showing the fusion junctions on mtDNA (MT) and the indicated CRISPR-Cas9 target sites (colorful triangles) on the nuclear DNA of HEK293T cells. The outer circle shows the human genome, labeled with numbers or characters. The colorful lines indicate the fusion between the target site and mtDNA. Annotations of mtDNA are shown at the bottom. D. Circos plot showing the fusion junctions on mtDNA (MT) and the indicated CRISPR-*Lb*Cas12a target sites (colorful triangles) on the nuclear DNA of HEK293T cells. Legends are described in C. E. Circos plot showing the fusion junctions on mtDNA (MT) and the indicated CRISPR-CasMINI target sites (colorful triangles) on the nuclear DNA of HEK293T cells. Legends are described in C. F. Box plot showing the frequency of mtDNA-nuclear DNA fusion events out of editing events at CRISPR-Cas target sites (colorful dots) under editing of *Sp*Cas9 variants. G. Distribution of mtDNA-nuclear DNA fusion caused by high fidelity Cas9 variants in the mES cells. Legends are described as depicted in C. H. Frequency of mtDNA-nuclear DNA fusion at *DNMT1*, *EMX1*, *MYC2*, and *RAG1_C* loci after editing by Cas9, BE4max, and ABEmax. EMX1 and MYC2 loci were not targetable by ABEmax. N.D., not determined.

High-fidelity *Sp*Cas9 variants such as eSpCas9, HF1, FeCas9, EvoCas9, HiFiCas9, Hypa, LZ3, and Sniper Cas9, as well as PAM-flexible *Sp*Cas9 variants including Cas9-NG, xCas9, SpG, and SpRY, have been developed to enhance editing specificity or broaden editing scope, respectively (Schmid-Burgk et al., 2020; Zhang et al., 2021). These *Sp*Cas9 variants exhibited comparable levels of mtDNA-nuclear DNA fusions to wild-type *Sp*Cas9 (Fig. 1F, Supplementary Table 3). Considering the inheritability of editing products in embryonic stem cells (ESCs), we also performed gene editing with five high-fidelity *Sp*Cas9 variants (Hypa, HF1, FeCas9, eCas9, and SuperFi) in parallel with *Sp*Cas9 in mouse ESCs (Zhang et al., 2021; Bravo et al., 2022). Consistently, we identified mtDNA integrations at the *c-Myc* target site in mouse ESCs (Fig. 1G). In contrast, by avoiding DSB generation, base editors significantly reduced mtDNA-nuclear fusion products at target loci (Fig. 1H) (Yin et al., 2022b). In total, we have identified over 1,400 mtDNA-nuclear DNA fusion events induced by various CRISPR-Cas editing tools. The mtDNA fusion junctions were distributed throughout the mitochondrial genome (Fig. S1E) and over 40% of mtDNA-nuclear DNA fusion events involved microhomology, primarily with 1-6 base pairs (Fig. S1F).

### Mitochondria-nuclear fusions undergo clonal expansion

The CRISPR-Cas9 system has been utilized to generate universal chimeric antigen receptor (CAR) T cells from human cord blood by disrupting the *TRAC*, *TRBC*, and *PDCD1* genes (Fig. 2A) (Yin et al., 2022b). The levels of mtDNA integrations in these edited primary immune cells were comparable to those observed in HEK293T cells, regardless of the position of target sites (Fig. S2A). Moreover, mtDNA integrations were detected in human CAR T cells after *ex vivo* culture for 3, 7, or 14 days (Fig. 2B and S2A) (Yin et al., 2022b). By labeling products with random molecular barcodes (RMB), PEM-seq enables the distinguishment of the same mtDNA-nuclear DNA fusion junction present in different expanded cells (Fig. 2A)(Wu et al., 2022). Notably, multiple chimeric junctions occurred in more than one cell (Fig. 2C), implying the expansion of mtDNA-nuclear DNA fusions during T cell culture *ex vivo*.

**Figure 2.**
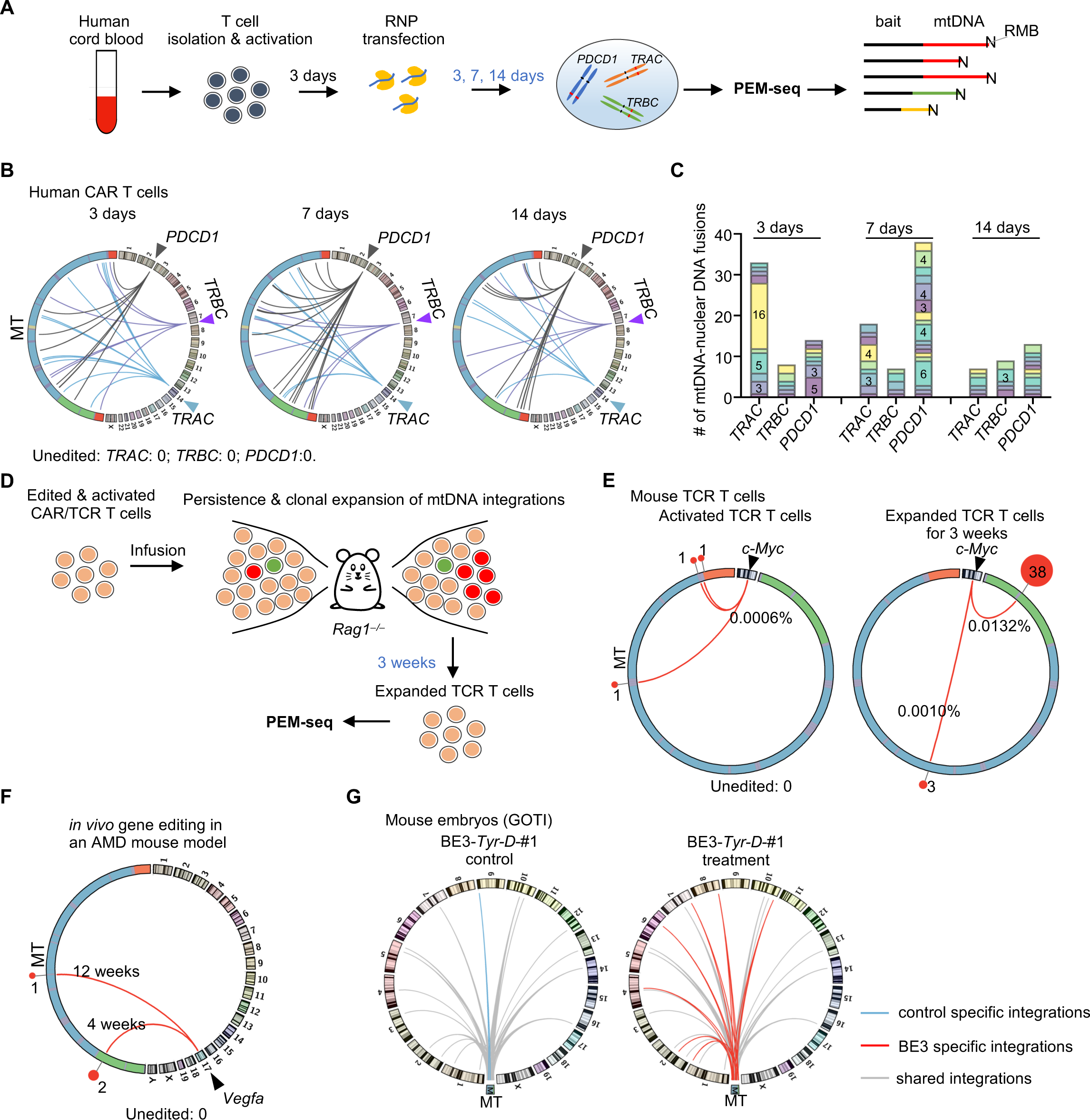
Mitochondrial DNA fuses to CRISPR-Cas9 target sites of the nuclear DNA *ex vivo* and *in vivo*. A. Schematic of ex vivo human T cell editing and PEM-seq. Briefly, human primary T cells isolated from cord blood were activated for 3 days and subsequently transfected with Cas9/gRNA ribonucleoprotein (RNP) complexes targeting *TRAC*, *TRBC*, and *PDCD1* loci. Cells were collected after 3, 7, or 14 days and subjected to PEM-seq libraries. In PEM-seq libraries, all products are labeled with random molecular barcodes (RMB) to distinguish biological expansion and PCR duplications. mtDNA-nuclear DNA fusions originate from the same mtDNA junction are labeled in red. B. Distribution of mtDNA-nuclear DNA fusions after CRISPR-Cas9 editing for 3, 7, and 14 days in human CAR T cells. Legends are described in Fig. 1B. C. Stacked bar plot showing the copy number of each mtDNA-nuclear DNA fusion type identified from *TRAC*, *TRBC*, and *PDCD1* bait after Cas9 editing for 3, 7, and 14 days. Numbers mark the copies of junctions with 3 copies or above. D. Schematic showing the amplification of mtDNA-nuclear DNA fusion through clonal expansion of TCR T cells after infusion. E. Distribution of mtDNA-nuclear DNA fusion in CRISPR-Cas9 treated mouse TCR-T cells before and post-infusion for 3 weeks. Legends are described as depicted in B. Percentages show the frequency of each mtDNA-nuclear DNA fusion out of Cas9-induced editing events. F. mtDNA-nuclear DNA fusions between *Vegfa* and mtDNA in an AMD mouse model treated with AAV containing CRISPR-Cas9. Legends are described as depicted in 1B. G. Distribution of mtDNA integration in the nuclear DNA post base editor (BE3) treatment in mouse embryos. Data were re-analyzed from GOTI libraries (SRA: SRP119022) (Zuo et al., 2019). The grey, blue, and red lines indicate integrations detected in both control and treatment, control-only, and treatment-only, respectively. MT, mtDNA.

To investigate the potential expansion of mtDNA integrations *in vivo*, we examined CRISPR-Cas9-edited mouse TCR T cells in a chronic inflammation murine model (Wu et al., 2022). Activated TCR T cells post Cas9 editing were infused into recipient mice for 3 weeks. Subsequently, both activated T cells and *in vivo* expanded T cells were subjected to PEM-seq analysis. Three different mtDNA-*c-Myc* fusions were identified in 167,611 editing events in activated T cells (Fig. 2E). Remarkably, one fusion junction was captured 38 times after infusion into the recipient mouse for 3 weeks (0.0132% of 286,818 editing events) (Fig. 2E). The identified 38 reads showed diverse RMBs and prey sequences, indicating that these fusions were originated from different cells but not PCR duplication (Fig. S2B). Moreover, mtDNA integrations were also detected in gene editing with an age-related macular degeneration (AMD) murine model *in vivo* (Fig. 2F), where adeno-associated virus (AAV) was used to target the *Vegfa* gene in retinal cells (Yin et al., 2022a). One mtDNA-*Vegfa* fusion event was found in 27,749 editing events from two different cells 4 weeks post-injection, and one fusion was captured in 35,534 editing events 12 weeks post-treatment (Fig. 2F). Collectively, these data suggest that mtDNA integrations can be amplified multiple times as the edited cells undergo clonal expansion.

The CRISPR-based base editors enable gene correction in the nuclear genome without directly inducing DSBs (Anzalone et al., 2020). To assess the nuclear base editors, the genome-wide off-target analysis by two-cell embryo injection (GOTI) assay was developed, which edited the embryos at the two-cell stage and collected cells from murine E14.5 embryos for whole-genome sequencing (Zuo et al., 2019). We employed the NUMTs-detection algorithm (Wei et al., 2022a) to analyze the discordant reads of GOTI libraries with base editor 3 (BE3) targeting the *Tyr-D* sites (Zuo et al., 2019). For the first set, we detected 14 shared mitochondrial segments in both control and BE3-edited cells. The BE3-edited cells had seven new mitochondrial segments and one missing segment compared to the control cells genome-wide (Fig. 2G), suggesting a potentially higher level of mtDNA integrations after base editing. Similar findings were obtained for the second set (Fig. S2C). Taken together, mtDNA integrations persisted during the development of mouse embryos.

### Mitochondrial perturbance enhances mtDNA transfer into nuclear DNA

Mitochondrial integrations into the nuclear target sites require the fusion of DNA breaks at both the mtDNA and nuclear DNA. To explore the contributions of mtDNA breaks to mitochondrial integrations during gene editing, we induced mitochondrial stress that can generate mtDNA breakage and release fragmented mtDNA into the cytosol (Xian et al., 2022; Kim et al., 2019). We employed carbonyl cyanidem-chlorophenyl hydrazone (CCCP) to disrupt the membrane potential and used paraquat to induce oxidative damage (O’Malley et al., 2020; Fessler et al., 2020; Riley et al., 2018). The levels of mtDNA integrations were found to be approximately 2-fold higher after treatment with CCCP or paraquat, as determined by PEM-seq (Fig. 3A and 3B). For further validation, we developed <u>inse</u>rtion-enriched target sequencing (Insert-seq), which employed two rounds of targeted PCR and two rounds of size selection to enrich fragment insertions at the nuclear editing site (Fig. 3C and S3A). Insert-seq mainly captured DNA fragment insertions less than 500 bp with a median length of 163 bp (Fig. S3A and S3B). We performed Insert-seq analysis on the same genomic DNA used for PEM-seq. The mtDNA integrations at the *c-MYC* target site accounted for approximately 0.43% and 0.42% in the presence of CCCP and paraquat, respectively, *via* Insert-seq. In contrast, the integration level was only 0.11% in the absence of mitochondrial stresses (Fig. 3D).

**Figure 3.**
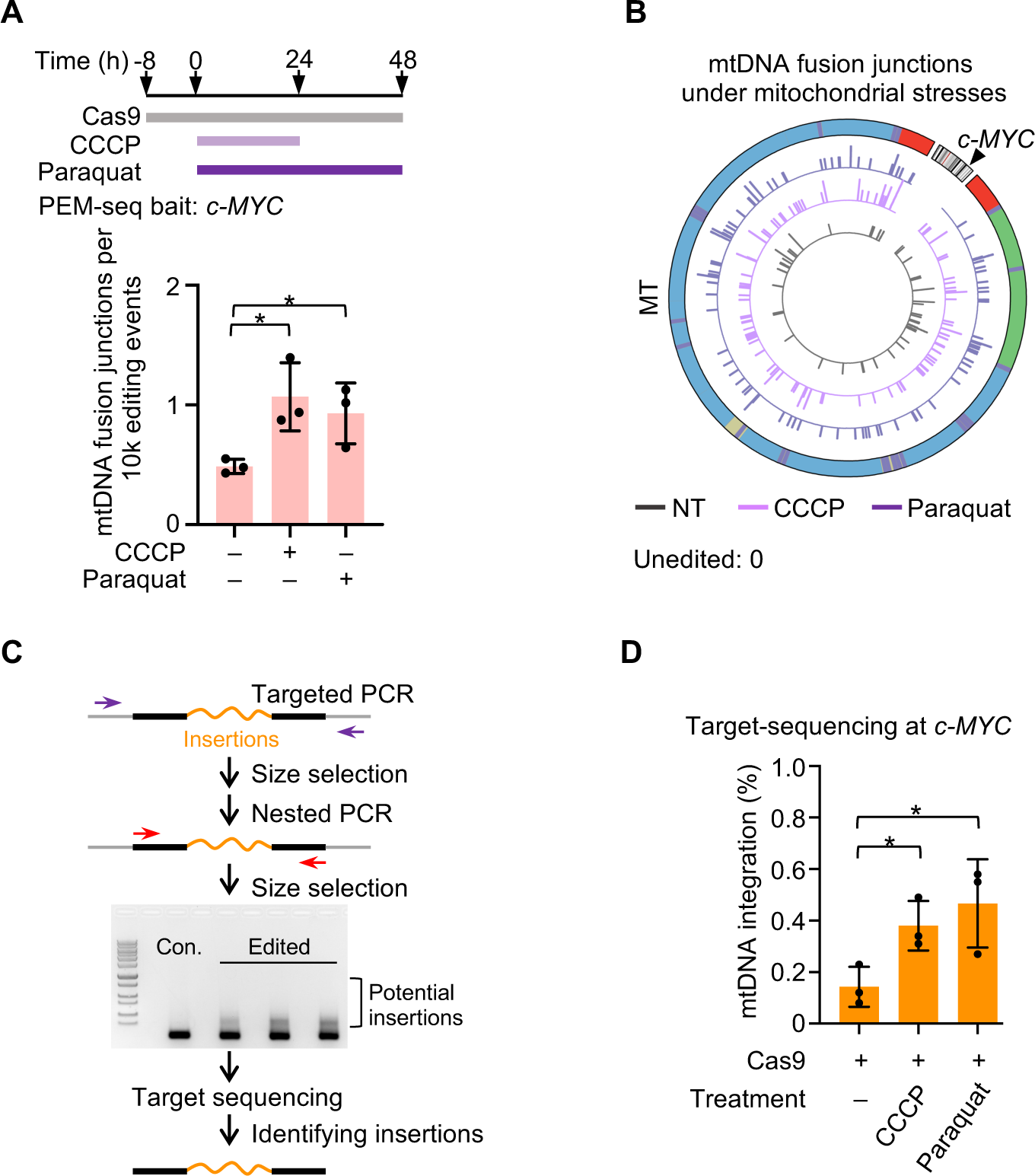
Mitochondrial stresses exacerbate mtDNA integration into the nuclear genome. A. Mitochondrial stresses inducing mtDNA-nuclear DNA fusions captured by PEM-seq. Top: cells were transfected with a plasmid containing *Sp*Cas9 and gRNA targeting *c-MYC* for 8 hours to allow the assembly of *Sp*Cas9 and gRNA. Subsequentially, cells are treated with CCCP or paraquat for 24 or 48 hours, respectively, and then harvested for PEM-seq and Insert-seq analysis. Bottom: percentages of mtDNA-nuclear DNA fusions captured by PEM-seq. Each dot represents a biological replicate. Mean±SD; *t*-test; *, *p*<0.05. B. Distribution of mtDNA (MT) fusion junctions captured by the *c-MYC* bait (black triangle and grey bars) with or without mitochondrial stresses (CCCP, light purple bars; paraquat, dark purple bars) treatment. Legends of mtDNA annotations are described as depicted in 1B. The inner circles show the number of each mtDNA-nuclear DNA fusion point on mtDNA in a log scale. C. Workflow of Insert-seq to enrich insertions (orange lines) at the Cas9 editing site (*c-MYC* locus). Briefly, two rounds of targeted PCR (purple and red arrows) are used to clone the editing events around the target site, followed by two rounds of size selection that enriches insertions. D. Percentages of mtDNA integrations within total insertions captured by Insert-seq at the *c-MYC* site in the presence or absence of mitochondrial stresses. Each dot represents a biological replicate. Mean±SD; *t*-test; *, *p*< 0.05; **, *p*< 0.01.

To investigate whether mtDNA breaks could be directly involved in nuclear DNA translocations, we employed mitoTALEN to generate DSBs in the *ND4* gene of the mitochondrial genome and performed PEM-seq by using the bait DSBs at the *c-MYC* locus on the nuclear DNA (Fig. 4A). We found that mitoTALEN treatment resulted in over 13-fold more mtDNA-nuclear DNA fusions than unedited cells (0.079% vs. 0.006% of total editing events) (Fig. 4B). The fusion junctions were highly enriched at the mitoTALEN target sites (Fig. 4C), as validated by PCR and sanger sequencing with two pairs of primers located adjacent to the editing sites of *c-MYC* and *ND4* (Fig. S4A and S4B). We detected many junctions over the entire mitochondrial genome beyond the *ND4* target site (Fig. 4C), consistent with previous findings that DSBs lead mainly to rapid fragmentation and degradation of the mtDNA (Silva-Pinheiro and Minczuk, 2022; Nissanka et al., 2018). We also conducted Insert-seq and identified over 6-fold more mitochondrial insertions in the presence of mitoTALEN than in untreated cells (0.39% vs. 0.06% of total insertions) (Fig. 4D and S4C), in line with PEM-seq. Of note, one mtDNA insertion at the *c-MYC* locus captured by Insert-seq harbored the same junctions as one insertion identified by PEM-seq (Fig. S4D). Collectively, these results suggest that fragmented mtDNA derived from mitochondrial stresses and DSBs can integrate into nuclear breaks.

**Figure 4.**
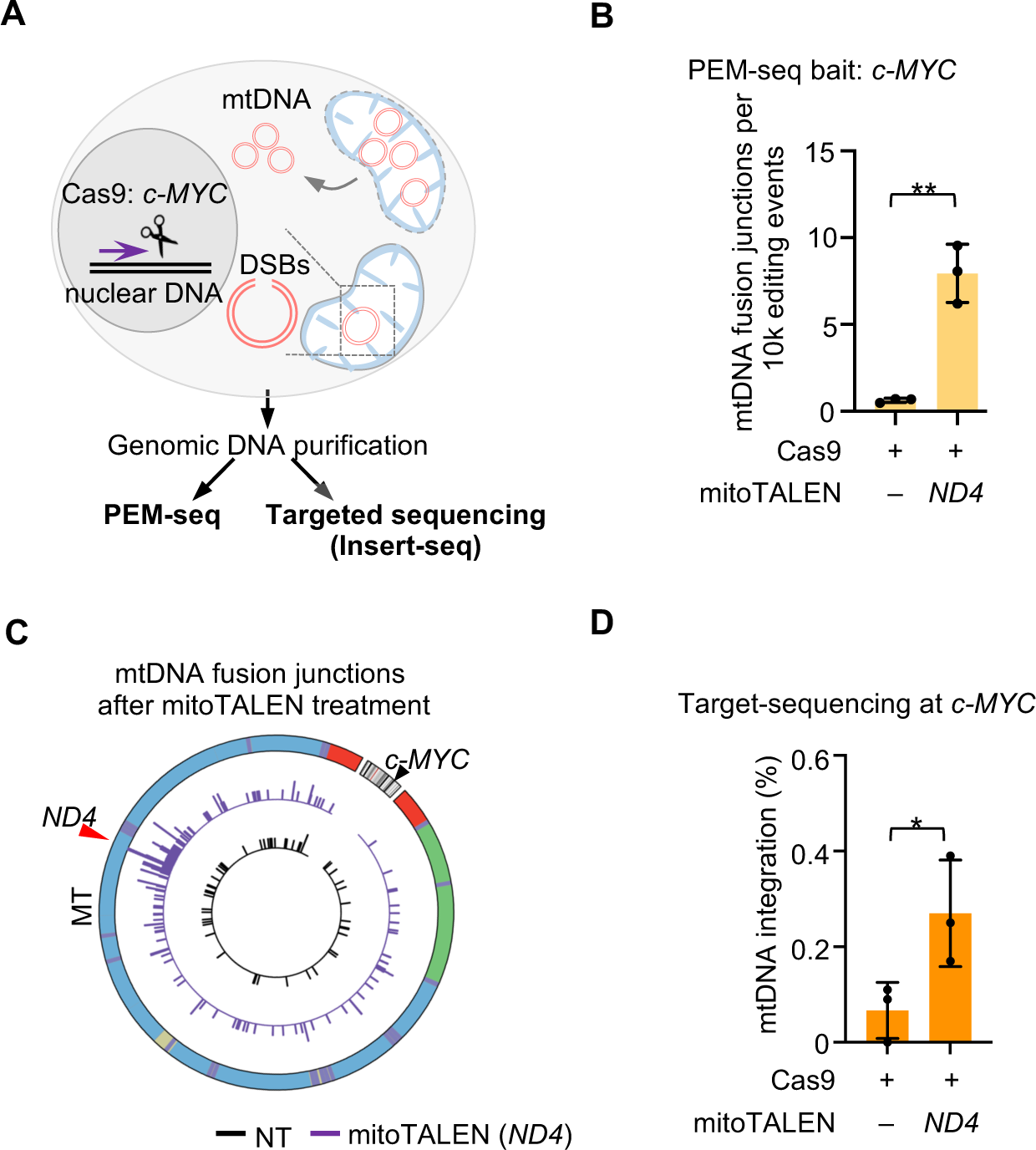
Mitochondrial editors exacerbate mtDNA integration into the nuclear target site. A. Illustration of mitochondrial stresses and mitoTALEN treatment induced mtDNA-nuclear DNA fusion captured by PEM-seq and Insert-seq. CRISPR-Cas9 (scissor) targets the *c-MYC* locus in nuclear DNA, and the primer (purple arrow) for PEM-seq is adjacent to the target site. B. Frequency of PEM-seq-captured mtDNA and the CRISPR-Cas9 target site fusion junctions with or without mitoTALEN treatment. Each dot represents a biological replicate. Mean±SD; *t*-test; **, *p*< 0.01. C. Distribution of mtDNA (MT) fusion junctions captured by the *c-MYC* bait (black triangle and grey bars) with or without mitoTALEN (*ND4* site, red triangle and purple bars) treatment. Legends are described as depicted in 3B. D. Percentages of mtDNA integrations within total insertions captured by Insert-seq at the *c-MYC* site. Each dot represents a biological replicate. Mean±SD; *t*-test; *, *p*< 0.05.

### Mitochondrial editors result in mtDNA integrations into the nuclear DNA

We next sought to investigate whether mitochondrial editing tools could induce the transfer of mtDNA into nuclear DNA. MitoTALEN has been developed to eliminate mitochondria carrying specific mutations and thus alleviate the mitochondrial heteroplasmy (Bacman et al., 2013; Bacman et al., 2018). MitoTALEN generated substantial DSBs within the editing window between the two TALE binding sites. To identify *bona fide* mitochondrial integrations, we placed the bait primer upstream of the left TALE binding site in the mitochondrial genome and only considered PEM-seq chimeric reads harboring mtDNA junctions within the editing window (Fig. 5A). We used mitoTALEN to edit the *ND5.1* gene and observed 7-fold more mtDNA-nuclear DNA fusions than in untreated cells (Fig. 5B and 5C). The identified nuclear junctions were evenly distributed on the chromosomes, with about 50.9% (57 of 112) of these junctions located in gene regions. Similarly, 5-fold more mtDNA-nuclear DNA fusions were captured in cells post mitoTALEN editing at the *ND4* gene (Fig. S5A and S5B).

**Figure 5.**
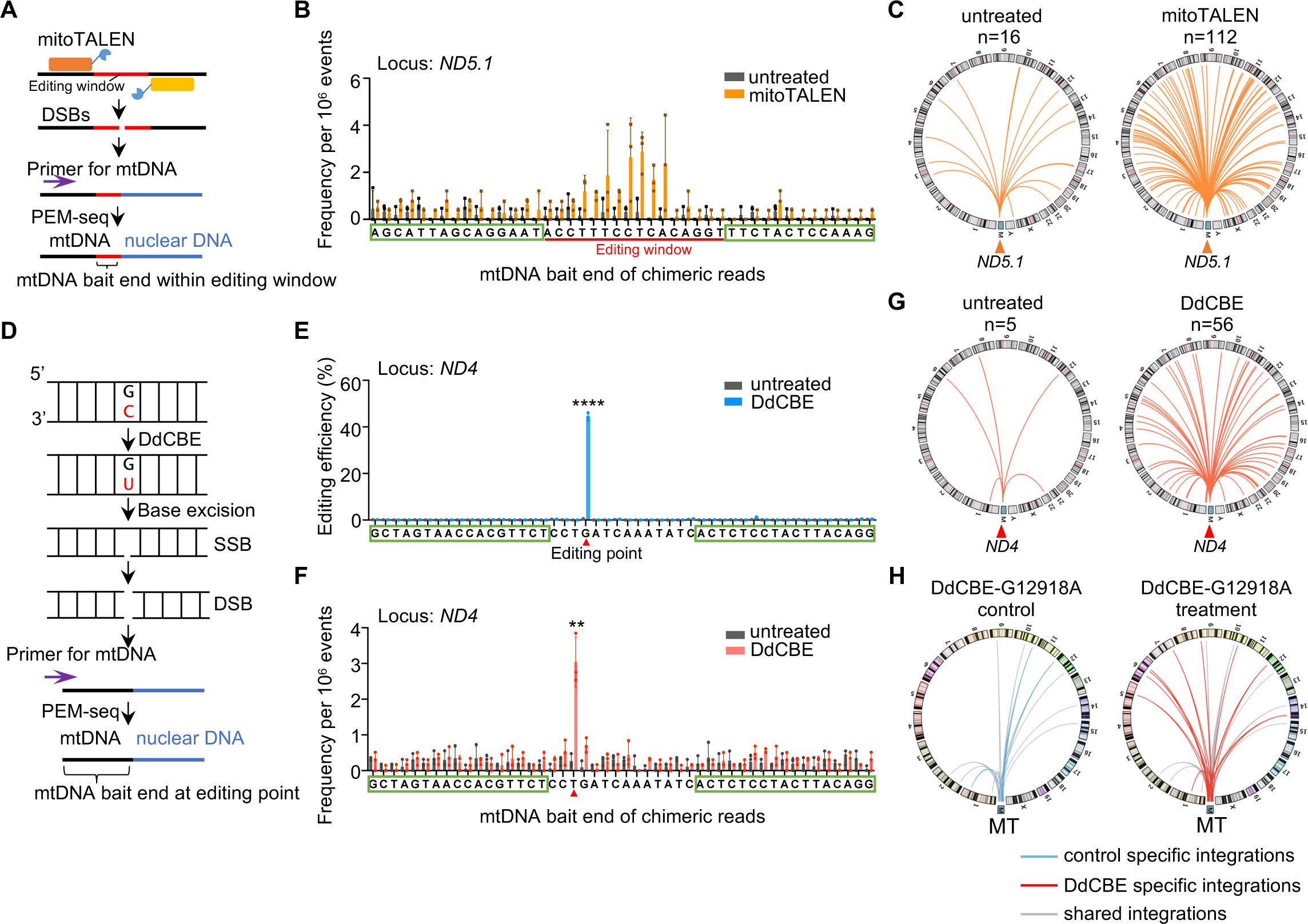
Mitochondrial editors result in mtDNA integrations into nuclear DNA. A. Schematic of mitoTALEN treatment induced mtDNA integration into the nuclear DNA, which can be captured by a primer (purple arrow) located adjacent to the mitoTALEN editing window (red line). mitoTALEN generated DSBs within the editing window between two TALE target sites. Chimeric reads containing *bona fide* mitoTALEN-induced mtDNA-nuclear DNA fusion should harbor mtDNA bait junctions within the editing window. B. Frequency of mtDNA-nuclear DNA fusions with mtDNA bait junctions at each nucleotide spanning the mitoTALEN target site on *ND5.1* (green boxes), editing window of mitoTALEN is marked with a red line. Each dot represents a biological replicate. Mean±SD; *t*-test; *, *p*< 0.05; **, *p*< 0.01. C. Distribution of mtDNA-nuclear DNA fusions (orange lines) with mtDNA bait junctions within the editing window of mitoTALEN. D. Illustration of DdCBE caused mtDNA-nuclear DNA integration. Legends are described as depicted in A. Chimeric reads containing *bona fide* DdCBE-induced fusion should harbor mtDNA bait junctions at the editing site. E. Editing efficiency (C to T or G to A) of DdCBE on *ND4*. The green boxes and the red triangle indicate the binding site and editing point of DdCBE. Each dot represents a biological replicate; Mean±SD; *t*-test; ****, *p*< 0.0001. F. Frequency of mtDNA-nuclear DNA fusions with mtDNA bait junctions at each nucleotide spanning the DdCBE target sites (green boxes) on ND4. Each dot represents a biological replicate. Mean±SD; *t*-test; **, *p*< 0.01. G. Distribution of mtDNA-nuclear DNA fusions (orange lines) with mtDNA ending at the editing site of DdCBE. H. Distribution of mtDNA-nuclear DNA fusions posts DdCBE treatment in mouse embryos. Data were re-analyzed from GOTI libraries (SRA: PRJNA786071) (Wei et al., 2022b). Legends are described as depicted in 2G. The grey, blue, and red lines indicate fusions detected in both control and treatment, control-only, and treatment-only, respectively.

The DdCBE enables CRISPR-free nucleotide modification in mtDNA, providing an important tool for mtDNA manipulation and disease therapy (Mok et al., 2020). We next investigated whether DdCBE could induce mtDNA integrations by cloning from *ND4* or *ND5.3* target site (Fig. 5D). At the *ND4* target site, the editing efficiency exceeded 40% (Fig. 5E). To identify *bona fide* mtDNA integrations, we only counted the chimeric junctions containing G to A mutation or missing the G nucleotide due to excision repair (Fig. 5D, 5F, and S5C). Post DdCBE treatment, we observed an 11-fold increase in mtDNA-nuclear DNA fusions at the *ND4* locus compared to untreated cells (Fig. 5F and 5G). Similarly, 4.6-fold more fusions were observed when DdCBE targeted the *ND5.3* locus (Fig. S5D-F). Besides, we also analyzed published GOTI data with DdCBE to induce the G12918A or C12336T mutation (Wei et al., 2022b). For G12918A editing, we detected eight shared mitochondrial segments in both DdCBE-edited and untreated cells, with the DdCBE-edited cells having 10 new mitochondrial segments and three missing segments compared to the untreated cells (Fig. 5H). Similar findings were obtained for C12336T editing (Fig. S5G). Altogether, these data indicated that DdCBE-editing can induce the transfer of mtDNA into the nuclear DNA.

### TREX1 and TREX2 suppress the transfer of mtDNA into nuclear DNA

The transfer of mtDNA into the nuclear DNA necessitates mtDNA breakage, and thereby it is conceivable that ectopic expression of exonucleases might promote the degradation of damaged mtDNA and prevent mtDNA integration into the nuclear DNA (Fig. 6A). TREX1 and TREX2 are efficient at degrading free cytosol DNA and have been applied in nuclear gene editing (Yin et al., 2022b). We generated TREX1n by removing the C-terminal cellular trafficking domain from TREX1, which did not affect its exonuclease activity. We fused TREX1n and TREX2 with DdCBE directly or with a mitochondrial targeting sequence (MTS) to co-express with DdCBE (Fig. 6A). The fusion and co-expression of TREX1n/TREX2 had no significant impact on the editing efficiency of DdCBE or the mtDNA copy numbers (Fig. 6B and 6C). We used these mitochondrial editing tools to edit the *ND4* gene in HEK293T cells and detected 12 mtDNA-nuclear DNA fusions after DdCBE editing *via* PEM-seq. In comparison, five to seven fusions were identified when cells were treated with DdCBE either fused or co-expressed with TREX1n/TREX2 (Fig. 6D), implying a potentially lower transfer rate of mtDNA into the nuclear DNA.

**Figure 6.**
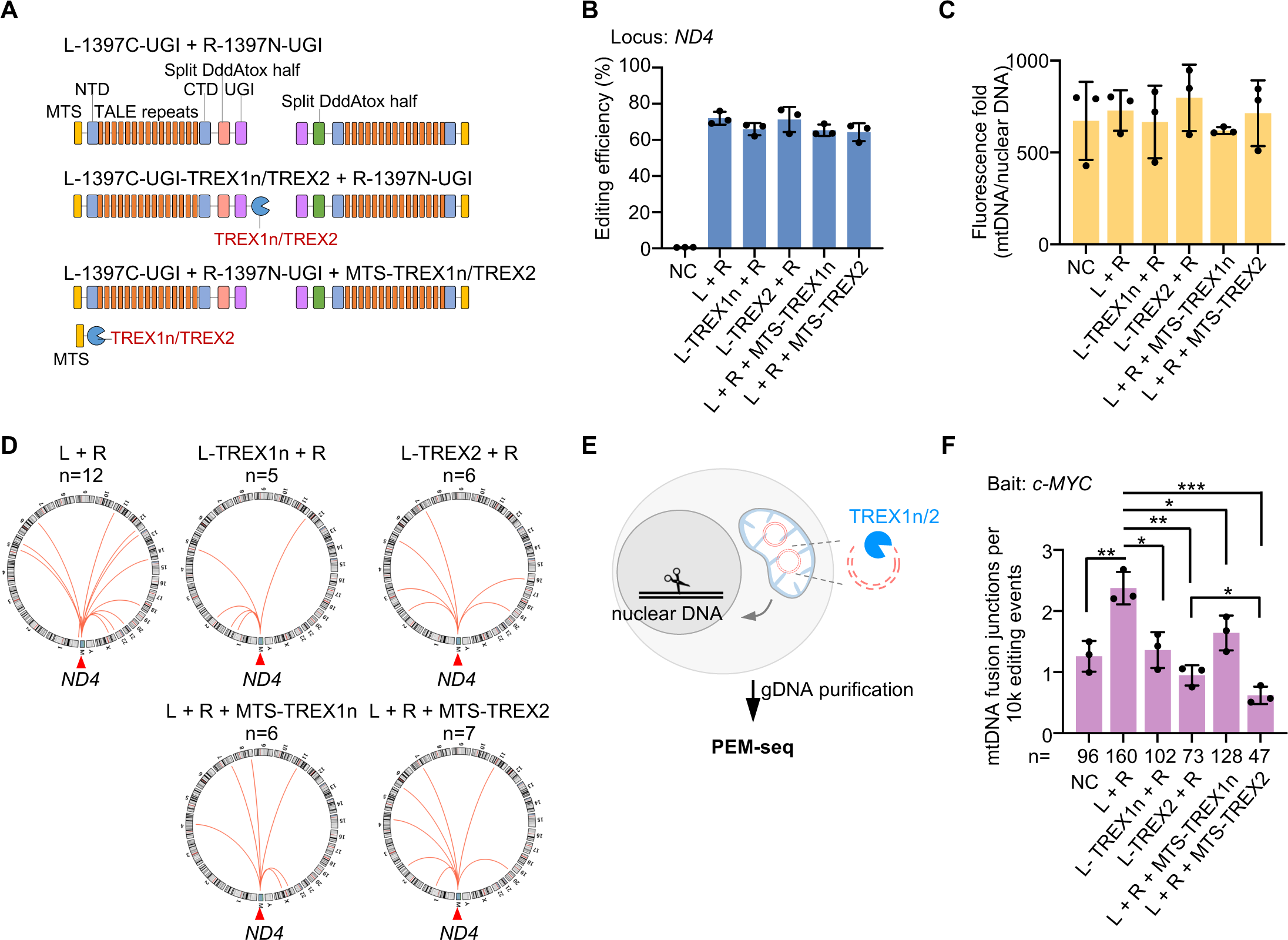
TREX1 and TREX2 suppress the transfer of mtDNA into nuclear DNA. A. Structures of DdCBE with or without TREX1n/TREX2. For the fusion form, TREX1n or TREX2 was fused to the C-terminal domain of L-1397C-UGI. Regarding separated TREX1n or TREX2, both nucleases were tagged with mitochondrial targeting sequence (MTS) on the N-terminal. L-1397C-UGI, left TALE arrays fused to C-terminal DddAtox half and UGI; R-1397N-UGI, right TALE arrays fused to N-terminal DddAtox half and UGI; NTD, N-terminal domain; CTD, C-terminal domain. B. Editing efficiency of DdCBE with or without TREX1n/TREX2 treatment. NC, negative control. Mean±SD. L, L-1397C-UGI; R, R-1397N-UGI. C. Fluorescence fold change of mtDNA to nuclear DNA detected by qPCR. D. Distribution of mtDNA-nuclear DNA fusions (orange lines) with mtDNA bait junctions ending at the editing site of DdCBE on *ND4*. The number (n) of fusions in each sample is normalized to the same editing events. L, L-1397C-UGI; R, R-1397N-UGI. E. Illustration of PEM-seq cloning from the nuclear DNA to assess DdCBE-induced mtDNA-nuclear DNA fusion with or without TREX1n/TREX2 treatment. F. Frequency of DdCBE-induced mtDNA fusing with the CRISPR-Cas9 target site with or without TREX1n/TREX2 treatment. Each dot represents a biological replication. Mean±SD; *t*-test; *, *p*< 0.05; **, *p*< 0.01; ***, *p*< 0.001. L, L-1397C-UGI; R, R-1397N-UGI.

To improve the throughput of mtDNA-nuclear DNA fusions for better statistical analysis, we introduced recurrent DSBs at *c-MYC* in the nuclear genome by *Sp*Cas9 and used them as bait to capture DdCBE-induced mtDNA fragmentation *via* PEM-seq (Fig. 6E). The *c-MYC* bait successfully captured 47 to 160 mitochondrial integrations for each treatment after normalization (Fig. 6F). We found 2-fold more mtDNA-nuclear DNA fusions post DdCBE treatment at the *ND4* locus, consistent with our above findings. However, fusion or co-expression of DdCBE with TREX1n/TREX2 resulted in a significant decrease in mtDNA-nuclear DNA fusions, with levels dropping to those of untreated cells (Fig. 6F). Moreover, TREX2 showed a relatively lower frequency of mtDNA-nuclear DNA fusion than TREX1n, and the MTS-tagged TREX2 slightly outperformed DdCBE-TREX2 (Fig. 6F).

## Discussion

The NUMTs with fragmented or full-length mtDNA have been widely observed in somatic and cancer cells (Wei et al., 2022a; Wei et al., 2020; Ju et al., 2015; Dayama et al., 2014; Mishmar et al., 2004; Woischnik and Moraes, 2002; Hazkani-Covo et al., 2010; Yuan et al., 2020). *De novo* transfer of mtDNA to the nuclear genome can lead to altered gene expression, such as the overactivation of *c-MYC* and *KCNMA1* oncogenes that contribute to tumor progression (Shay et al., 1991; Ju et al., 2015), or the disruption of genes, which may contribute to the development of Pallister-Hall syndrome and mucolipidosis (Turner et al., 2003; Goldin et al., 2004; Borensztajn et al., 2002). In addition, mtDNA integrations lead to mitochondrial heteroplasmy, and the level of mitochondrial heteroplasmy in individuals could change rapidly during development (Schwartz and Vissing, 2002), similar to our findings that mtDNA integrations can be amplified during the clonal expansion of CAR or TCR T cells (Fig. 2A-E).

Both the nuclear and mitochondrial genomes encounter spontaneous DNA lesions triggered by oxidative stress, ultraviolet (UV) light, chemicals, and replicative stress (Tubbs and Nussenzweig, 2017). Furthermore, the mitochondrial genome undergoes fragmentation during stresses, resulting in the leakage of multiple mtDNA fragments into the cytosol and nucleus (Xian et al., 2022; Kim et al., 2019). In this study, we use the gene editing-induced DSBs to show that any DSB in the nuclear genome might capture these mtDNA fragments, not limited to DSBs in physiological cellular processes such as V(D)J recombination (Lebedin et al., 2022). In this context, a 41-bp mtDNA was found to be integrated at the junction of a reciprocal constitutional translocation (Willett-Brozick et al., 2001). Therefore, NUMTs can occur in different types of cells and may become inheritable if they occur in ES cells. In this context, hundreds of NUMTs have been embedded in the human genome. Since most cancer cells are susceptible to DNA damage and possess a high oxidative metabolism, *de novo* NUMTs are quite frequent in cancer cells (Wei et al., 2022a; Kopinski et al., 2021).

The CRISPR-free DdCBE system and other forms of mitochondrial base editors have demonstrated high efficiency in modifying mtDNA and offer great potential for treating mitochondria-related diseases (Mok et al., 2020; Mok et al., 2022; Cho et al., 2022). However, our findings reveal a previously unknown risk that both nuclear and mitochondrial editing systems, such as DdCBE, can cause the transfer of mtDNA into the nuclear genome and nearly half of these mitochondrial integrations occur in gene regions. In the context of 10^8^∼10^9^ edited CAR or TCR T cells being infused into patients during therapy (Stadtmauer et al., 2020), one in 10^4^∼10^5^ infused cells may carry deleterious mtDNA-nuclear DNA fusions. Therefore, caution should be exercised when using both nuclear and mitochondrial editors for therapeutic purposes. Fortunately, we have shown that the fusion or co-expression of DdCBE with TREX1/TREX2 can degrade mitochondrial fragments and prevent the fusion of mtDNA with DSBs in the nuclear genome. In support, TREX1 mutations found in autoimmune diseases were accompanied by increased amounts of escaped mtDNA fragments (Hemphill and Perrino, 2019; Skopelja-Gardner et al., 2022). In particular, TREX2, located in the mitochondria, outperforms DdCBE-fused TREX1n/TREX2 or co-expressed TREX1n, and has no impact on mtDNA copy numbers, providing a feasible approach to enhance the safety of mitochondrial base editors.

## Materials and Methods

### Cell culture and transfection

V6.5 mouse embryonic stem cells (a gift from Dr. Xiong Ji, School of Life Sciences, Peking University) were cultured in Knockout^TM^ Dulbecco′s Modified Eagle′s Medium (DMEM, Gibco) containing 15% fetal bovine serum (Gibco), MEM non-essential amino acids solution (Sigma), nucleotides (Millipore), penicillin-streptomycin (Gibco), L-glutamine solution (Sigma), 2-mercaptoethanol (Sigma), LIF (Millipore), CHIR99021 (Selleck), and PD0325901 (Selleck) at 37 °C under 5% CO_2_. One million cells were transfected with 2 μg plasmid containing puromycin resistance gene, gRNA, and Cas9 variants targeting the *c-MYC* locus by nucleofector (Lonza, 4D-Nucleofector X). The transfected cells were treated with 1 μg/mL puromycin for 1.5 days after one day post-transfection, and then cultured with fresh medium for another day. Genomic DNA was extracted for PEM-seq analysis.

HEK293T cells were cultured in DMEM (Gibco) supplemented with 10% fetal bovine serum (ExCell Bio), penicillin-streptomycin (Gibco), and L-glutamine solution (Sigma) at 37 °C with 5% CO_2_. HEK293T cells cultured in 10-cm dishes were transfected with *Sp*Cas9-gRNA plasmid targeting *c-MYC* locus (15 μg) by calcium phosphate, with or without additional mitoTALEN plasmid targeting the *ND4* locus (12 μg *ND4*-L1397C-LEFT+12 μg *ND4*-L1397C-RIGHT). Regarding the mitochondrial stress treatment, 15 μg plasmid expressing *Sp*Cas9 targeting *c-MYC* locus was transfected into HEK293T cells by calcium phosphate. The transfected cells were then treated with 10 μM CCCP for 1 day and then cultured with fresh medium for another day, or treated with 500 μM paraquat for 2 days. After that, cells were harvested and the genomic DNA was extracted for PEM-seq analysis.

To determine the occurrence of mtDNA-nuclear DNA fusion caused by mtDNA editors. HEK293T cells were transfected with mitoTALEN targeting *ND5.1* (20 μg *ND5.1*-L1397N-LEFT + 20 μg *ND5.1*-L1397N-RIGHT) in 10-cm dishes for 3 days. Regarding DdCBEs, HEK293T cells were also transfected with DdCBE plasmids targeting *ND4* locus (20 μg *ND4*-L1397C-LEFT + 20 μg *ND4*-L1397C-RIGHT) or *ND5.3* locus (20 μg *ND5.3*-L1397C-LEFT + 20 μg *ND5.3*-L1397C-RIGHT) for 3 days. After transfection, cells were harvested and permeabilized with NP-40 buffer (10 mM HEPES-NaOH, pH 7.6; 10 mM KCl; 0.1 mM EDTA; 0.3% NP-40) for 10 min, followed by centrifugation for nuclei collection and genomic DNA extraction.

For the TREX1/2 co-expression analysis, HEK293T cells were transfected with *Sp*Cas9 plasmid targeting the *c-MYC* locus (15 μg) and DdCBE-*ND4*-L1397C plasmids (20 μg + 20 μg) containing TREX1n/TREX2 at the C-terminus of the UGI of the left arm of DdCBE. Besides, DdCBE-*ND4*-L1397C plasmids and MTS-tagged TREX1n/TREX2 driven by a separate promoter were also introduced into HEK293T cells. All sample cells were collected at 3 days post-transfection.

### PEM-seq library preparation and analysis

The PEM-seq libraries were prepared according to the previously reported procedures(Liu et al., 2022). Briefly, 50 μg of genomic or nuclear DNA from HEK293T cells, or 20 μg of genomic DNA from mES cells was generally required for each library. PEM-seq libraries were sequenced on Illumina Hiseq platforms, 2×150 bp. Primers used in this study were listed in the Key Resource Table.

The PEM-seq data was analyzed using the PEM-Q pipeline and aligned to the hg38 genome assembly as described(Liu et al., 2022). To identify mtDNA fusions, we only kept chimeric reads mapped to “chrM” with a MAPQ >30. Similarly, when analyzing PEM-seq data cloned from the mtDNA, we only kept chimeric reads mapped to the nuclear DNA with a MAPQ >30 and then removed those reads mapped to repetitive genomic regions annotated in the UCSC database.

### Insertion-enriched target sequencing and analysis

Genomic DNA (gDNA) from cells with mitochondrial stresses or mitoTALEN treatment were subjected to both PEM-seq and insertion-enriched target sequencing analysis. Regarding insertion-enriched target sequencing, 2 μg gDNA was amplified by two primers (sequences were listed in the Key Resource Table) flanking the CRISPR-Cas9 target site and Taq DNA polymerase in 100 μL PCR mixture and then performed as follows: 95℃, 5 min; 95℃, 30 second, 59℃, 30 seconds, 72℃, 4 min, 20 cycles; 72℃, 5 min. DNA products were size selected by AMpure beads to keep insertions. After that, another two primers (sequences were listed in the Key Resource Table) were used to further enrich DNA products on the Cas9-editing site by nested PCR, and the program was set as follows: 95℃, 3 min; 95℃, 30 seconds, 59℃, 30 second, 72℃, 3 min, 15 cycles; 72℃, 5 min. A second size selection was performed to keep insertions by AMpure beads and the recovered DNA was tagged with Illumina adapter sequences. Library DNA was sequenced on Hi-seq platforms, 2×150 bp.

Raw reads were aligned to the human reference genome, assembly hg38, by BWA-MEM with the default parameters. The chimera reads, with MAPQ>30 for each segment, were kept for insertion identification. Reads beginning at the primer start sites, covering at least 40 bp of both upstream and downstream of the Cas9-editing sites, and containing insertion fragments of less than 2000 bp were potential insertions after genome editing. Insertions with ≥4 counts were kept for further analysis.

## Data availability

The PEM-Q pipeline is available at https://github.com/JiazhiHuLab.

## Contributions

J.H. conceived and supervised the project. J.W., Y.L., and J.H. designed the experiments; J.W., Y.L., L.O., T.G., J.L., S.Y., C.X., and J.Y. performed the experiments; J.W., Y.L., Z.Z., X.L., M.L., and J.H. analyzed the data; and J.W., Y.L., Y.T., and J.H. wrote the paper.

## Acknowledgments

We acknowledge all members of the Hu laboratory for their critical comments and helpful discussions. DdCBE plasmids were kind gifts from Dr. Chengqi Yi at Peking University. We also thank the Flow Cytometry Core at the National Center for Protein Sciences, Peking University, for the technical help.

## Supplementary Figure legends

**Figure S1.**
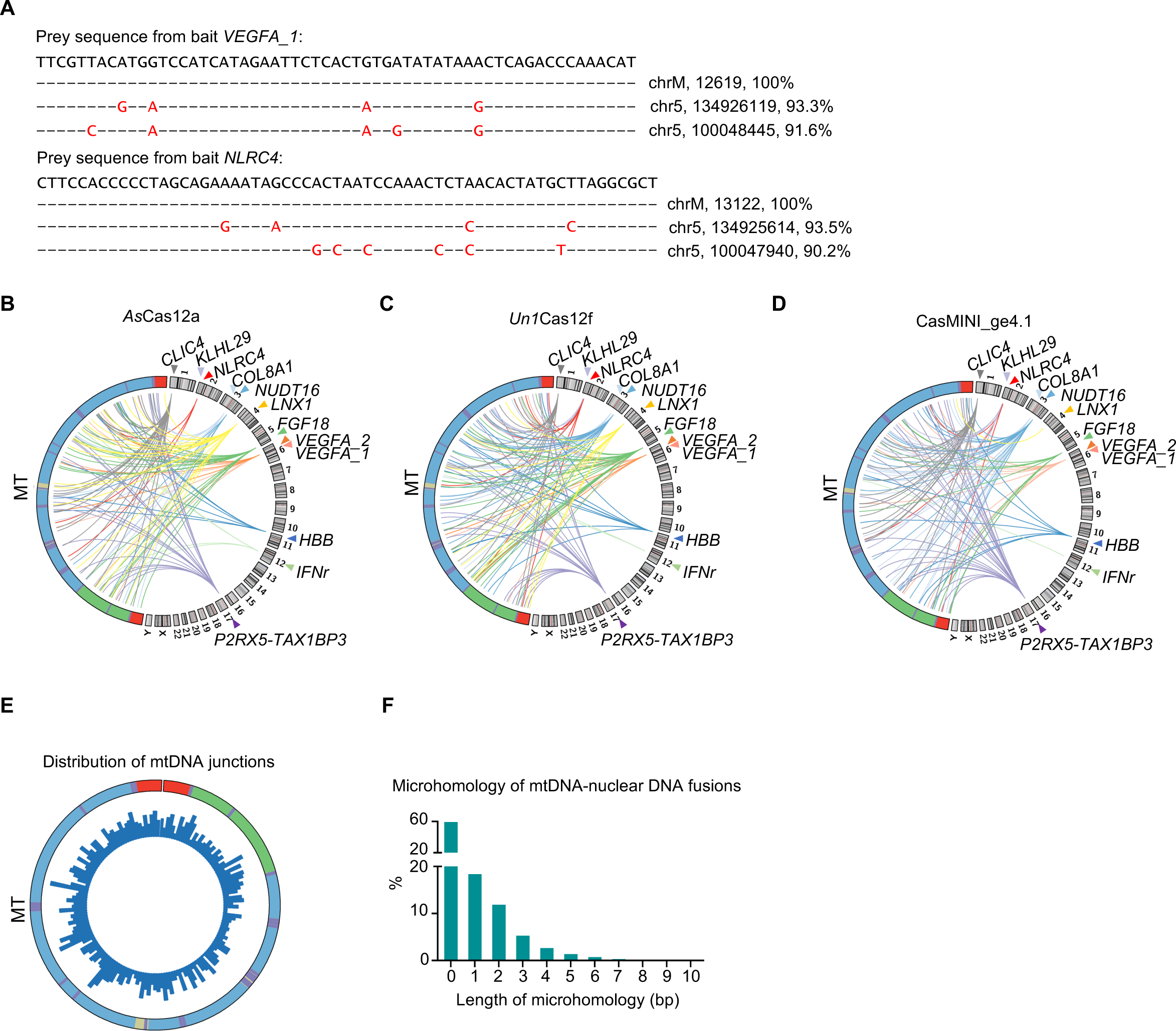
Identification of mtDNA fusion to nuclear targeting site of CRISPR-Cas systems. A. Representative mtDNA integration sequences that are aligned to the mitochondrial genome. B. Circos plot showing the fusion junctions on mtDNA (MT) and the indicated *Lb*Cas12a target sites (colorful triangles) on the nuclear DNA of HEK293T cells. Legends are described in 1B. C. Circos plot showing the fusion junctions on mtDNA (MT) and the indicated *Un1*Cas12f, target sites (colorful triangles) on the nuclear DNA of HEK293T cells. Legends are described in 1B. D. Circos plot showing the fusion junctions on mtDNA (MT) and the indicated CasMINI_ge4.1 target sites (colorful triangles) on the nuclear DNA of HEK293T cells. Legends are described in 1B. E. Distribution of mtDNA fusion points identified from CRISPR-Cas systems (1E and 1F). F. Percentage and length of microhomology identified in chimeric reads containing mtDNA-nuclear DNA fusion.

**Figure S2.**
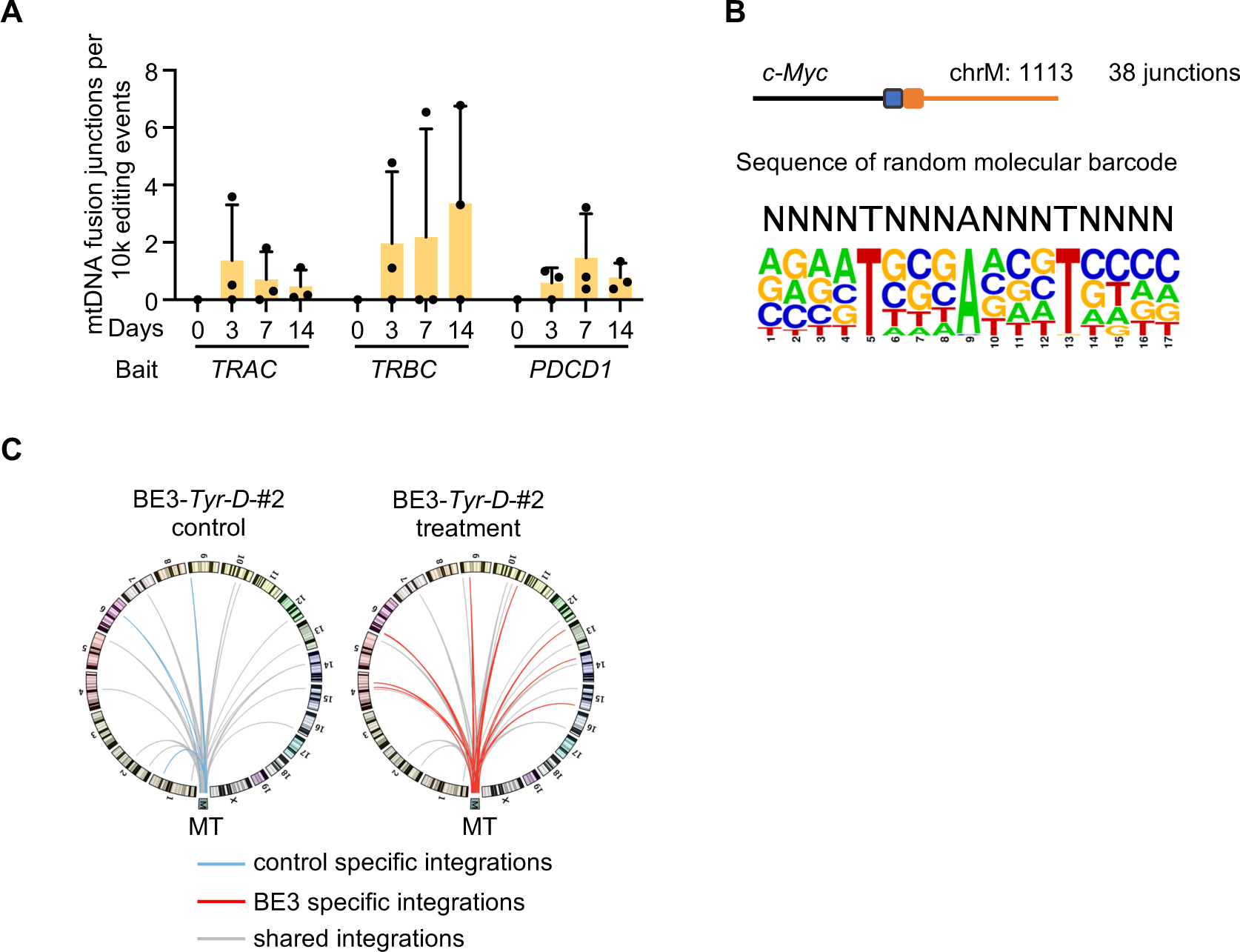
Mitochondrial DNA fuses to CRISPR-Cas9 targeting sites of the nuclear DNA *ex vivo* and *in vivo*. A. Percentages of mtDNA-nuclear DNA fusions at *TRAC*, *TRBC*, and *PDCD1* in human CAR-T cells, related to 2B. B. Sequence logo showing the frequency of nucleotides of random molecular barcodes from reads in 38 copies of mtDNA-nuclear DNA fusion in infused mouse TCR T cells at a single junction indicated in 2E. C. Distribution of mtDNA-nuclear DNA fusions posts base editor treatment in mouse embryos. Data were re-analyzed from GOTI libraries (SRA: SRP119022) (Zuo et al., 2019). Legends are described as depicted in 2G.

**Figure S3.**
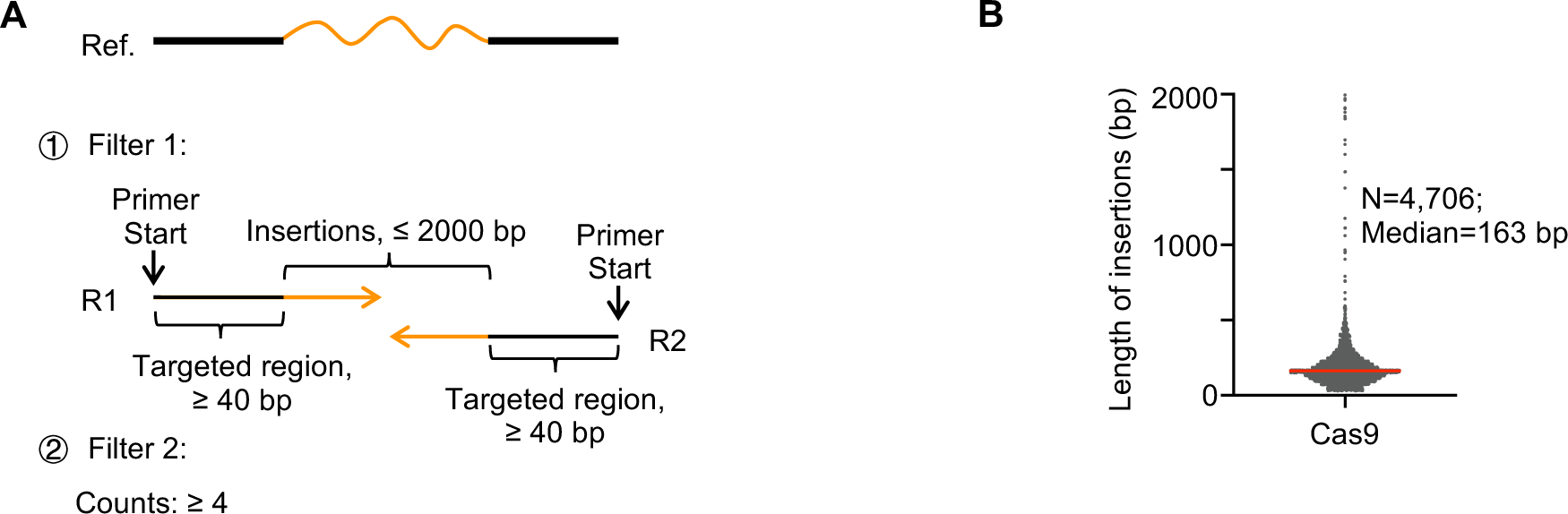
Using target sequencing for validation of mtDNA integrations. A. Schematic of Insert-seq captured *bona fide* insertions (orange lines) at the Cas9 editing sites. Briefly, reads should exactly start at the primer target site and then extend at least 40 bp on the target genome. The length of insertion should be less than 2000 bp, with a MAPQ >30. Moreover, each insertion must have more than 4 hits. B. Length distribution of Insert-seq captured insertions. Red line, median length.

**Fig. S4.**
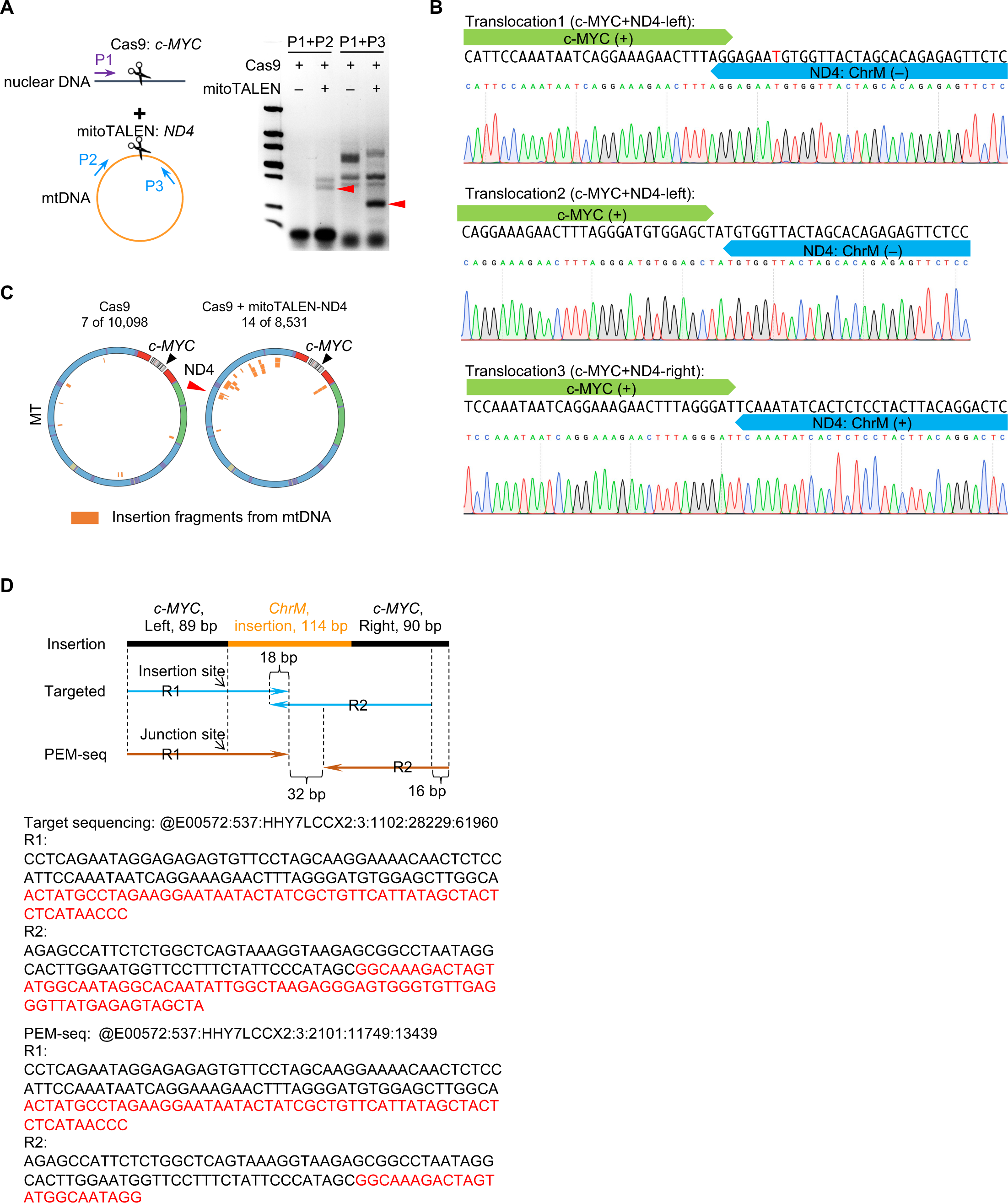
Mitochondrial DNA breaks exacerbate mtDNA integration into nuclear genome. A. PCR products showing the mtDNA-nuclear DNA fusion induced by Cas9 editing on *c-MYC* and mitoTALEN targeting *ND4.* Primers targeting nuclear DNA (purple arrow) and mtDNA (blue arrows) are used, and the expected products are labeled by red triangles. B. Sanger sequencing showing the fusion between *c-MYC* (green boxes) and *ND4* (blue boxes). C. Distribution of Insert-seq captured mtDNA segments that were inserted into *c-MYC*. D. Illustration of one mtDNA insertion that was captured by both PEM-seq and Insert-seq. Reads (R1 or R2) from PEM-seq (brown) and Insert-seq (blue) harbor the same insertion sites. Moreover, reads from Insert-seq shared an 18-bp overlap, while a 32-bp gap was found between R1 and R2 from PEM-seq. The sequences are listed below with mtDNA sequences marked in red.

**Figure S5.**
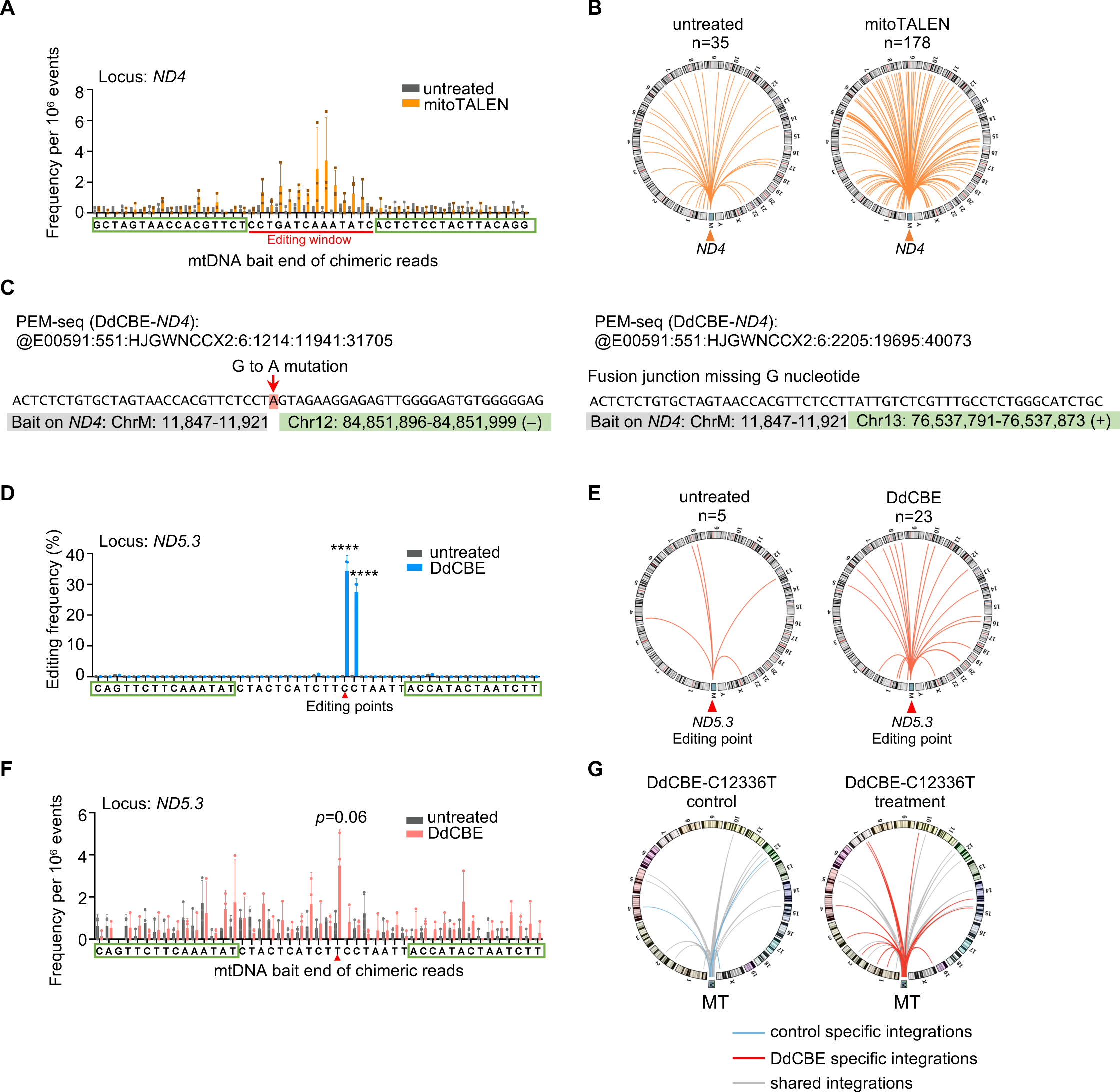
Mitochondrial editors result in mtDNA integrations into nuclear DNA. A. Frequency of mtDNA-nuclear DNA fusions with mtDNA bait junctions at each nucleotide spanning the mitoTALEN target site on *ND4*. Each dot represents a biological replicate. Mean±SD; *t*-test. B. Distribution of mtDNA-nuclear DNA fusions (orange lines) with mtDNA ending at the editing site of DdCBE. Legends are described as depicted in 5G. C. Representative sequence of *bona fide* DdCBE-induced mtDNA-nuclear DNA fusion. The red arrow marks the DdCBE caused G to A mutation. Light grey and green boxes mark the sequence aligned to mitochondrial and nuclear DNA. D. Editing efficiency of DdCBE on *ND5.3*. Three biological replicates; Mean±SD; *t*-test; ****, *p*< 0.0001. E. Distribution of mtDNA-nuclear DNA fusions (orange lines) with mtDNA bait junctions at the editing site of DdCBE. F. Frequency of mtDNA-nuclear DNA fusions with mtDNA bait junctions at each nucleotide spanning the DdCBE target site on *ND5.3*. Each dot represents a biological replicate. Mean±SD; *t*-test. G. Distribution of mtDNA-nuclear DNA fusion post mitochondrial base editor treatment in mouse embryos (SRA: PRJNA786071) (Wei et al., 2022b). Legends are described as depicted in 5H.

